# Efficient strategies for calculating blockwise likelihoods under the coalescent

**DOI:** 10.1101/016469

**Authors:** Konrad Lohse, Martin Chmelik, Simon H. Martin, Nicholas H. Barton

**Affiliations:** Institute of Evolutionary Biology University of Edinburgh King’s Buildings Charlotte Auerbach Road Edinburgh EH9 3FL, UK; Institute of Science and Technology Am Campus 1 A-3400 Klosterneuburg Austria; Zoology Department University of Cambridge UK

**Keywords:** Maximum likelihood, population divergence, gene flow, structured coalescent, generating function

## Abstract

The inference of demographic history from genome data is hindered by a lack of efficient computational approaches. In particular, it has proven difficult to exploit the information contained in the distribution of genealogies across the genome. We have previously shown that the generating function (GF) of genealogies can be used to analytically compute likelihoods of demographic models from configurations of mutations in short sequence blocks (Lohse *et al.,* 2011). Although the GF has a simple, recursive form, the size of such likelihood computations explodes quickly with the number of individuals and applications of this framework have so far been limited to small samples (pairs and triplets) for which the GF can be written down by hand. Here we investigate several strategies for exploiting the inherent symmetries of the coalescent. In particular, we show that the GF of genealogies can be decomposed into a set of equivalence classes which allows likelihood calculations from non-trivial samples. Using this strategy, we used *Mathematica* to automate block-wise likelihood calculations based on the GF for a very general set of demographic scenarios that may involve population size changes, continuous migration, discrete divergence and admixture between multiple populations. To give a concrete example, we calculate the likelihood for a model of isolation with migration (IM), assuming two diploid samples without phase and outgroup information, and compare the power of our approach to that of minimal pairwise samples. We demonstrate the new inference scheme with an analysis of two individual butterfly genomes from the sister species *Heliconius melpomene rosina* and *Heliconius cyndo.*

---

Genomes contain a wealth of information about the demographic and selective history of populations. However, efficiently extracting this information to fit explicit models of population history remains a considerable computational challenge. It is currently not feasible to base demographic inference on a complete description of the ancestral process of coalescence and recombination, and so inference methods generally rely on making simplifying assumptions about recombination (but see Rasmussen *et al.,* 2014). In the most extreme case of methods based on the site frequency spectrum (SFS), information contained in the physical linkage of sites is ignored altogether (Gutenkunst *et al.,* 2009; Excoffier *et al.,* 2013). Because the SFS is a function only of the expected length of genealogical branches (Griffiths & Tavaré, 1998; Chen, 2012), this greatly simplifies likelihood computations. However, it also sacrifices much of the information about past demography. Other methods approximate recombination along the genome as a Markov process (Li & Durbin, 2011; Harris & Nielsen, 2013). However, this approach is computationally intensive, limited to simple models (Schiffels & Durbin, 2014) and/or pairwise samples (Li & Durbin, 2011; Mailund *et al.,* 2012) and requires phase information and well assembled genomes which are still only available for a handful of species.

A different class of methods assumes that recombination can be ignored within sufficiently short blocks of sequence (Hey & Nielsen, 2004; Yang, 2002). The benefit of this “multi-locus assumption” is that it gives a tractable framework for analysing linked sites, and so captures the information contained in the distribution of genealogical branches. Multi-locus methods are also attractive in practice because they naturally apply to RAD data or partially assembled genomes that can now be generated for any species (e.g. Davey & Blaxter, 2011; Hearn *et al.,* 2014).

For small samples, the probability of seeing a particular configuration of mutations at a locus can be obtained analytically. For example, Wilkinson-Herbots (2008) and Wang & Hey (2010) have derived the distribution of pairwise differences under a model of isolation with migration (IM) and Wilkinson-Herbots (2012) has extended this to a history where migration is limited to an initial period. Yang (2002) derives the probability of mutational configurations under a divergence model for three populations and a single sample from each and Zhu & Yang (2012) have included migration between the most recently diverged pair of populations in this model. However, all of these particular cases can be calculated using a general procedure based on the generating function for the genealogy (Lohse *et al.,* 2011). Here, we explain how the GF, and – from it – model likelihoods can be efficiently computed for larger samples than has hitherto been possible.

## The generating function of genealogies

Assuming an infinite sites mutation model and an outgroup to polarize mutations, the information in a non-recombining block of sequence can be summarized as a vector *<u>k</u>* of counts of mutations on all possible genealogical branches *<u>t</u>.* Both *<u>t</u>* and *<u>k</u>* are labelled by the individuals that descend from them. We have previously shown that the probability of seeing a particular configuration of mutations *<u>k</u>* can be calculated directly from the Laplace Transform or generating function (GF) of genealogical branches (Lohse *et al.,* 2011). Given a large sample of unlinked blocks, this gives a framework for computing likelihoods under any demographic model and sampling scheme. Full details are given in Lohse *et al.* (2011). Briefly, the GF is defined as *ψ*[*<u>ω</u>* = *Ε*[e^−*<u>ω</u>*.<u>t</u>^], where *<u>ω</u>* is a vector of dummy variables corresponding to *<u>t</u>.* Setting the *<u>ω</u>* to zero necessarily gives one, the total probability; differentiating with respect to *ω_i_* and setting the *<u>ω</u>* to zero gives (minus) the expected coalescence time. If we assume an infinite sites mutation model, the probability of seeing *k_s_* mutations on a particular branch s is (Lohse *et al.,* 2011, eq. 1):

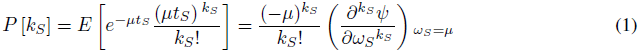

This calculation extends to the joint probability of mutations *P*[*<u>k</u>*]. Using the GF rather than the distribution of branches itself to compute *P*[*<u>k</u>*] is convenient because we avoid the Felsenstein (1988) integral and because the GF has a very simple form: going backwards in time, the GF is a recursion over successive events in the history of the sample (Lohse *et al.,* 2011, eq. 4):

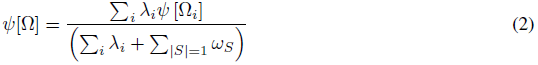
where *Ω* denotes the sampling configuration (i.e. the location and state of lineages) before some event *i* and *Ω_i_* the sampling configuration afterwards. Events during this interval occur with a total rate Σ*_i_* λ*_i_*. The numerator is a sum over all the possible events *i* each weighted by its rate λ*_i_*. Equation 2 applies to any history that consist of independently occuring events. As outlined by Lohse *et al.* (2011), the GF for models involving discrete events (population splits, bottlenecks) can be found by inverting the GF of the analogous continuous model. In other words, if we know the GF for a model that assumes an exponential rate of events at rate Λ, then taking the inverse LT wrt Λ gives the GF for any fixed time of the event.

In principle, the GF recursion applies to any sample size and model and can be automated using symbolic software (such as *Mathematica).* In practice however, likelihood calculations based on the GF have so far been limited to pairs and triplets: Lohse *et al.* (2011) computed likelihoods for an IM model with unidirectional migration for three sampled genomes and Lohse *et al.* (2012) and Hearn *et al.* (2014) derived likelihoods for a range of divergence histories for a single genome from each of three populations with instantaneous admixture, including the model used by Green *et al.* (2010) to infer Neandertal admixture into modern humans (Lohse & Frantz, 2014).

There are several serious challenges in applying the GF framework to larger samples of individuals. First, the number of sample configurations (and hence GF equations) grows super-exponentially with sample size. Thus, the task of solving the GF and differentiating it to tabulate probabilities for all possible mutational configurations quickly becomes computationally prohibitive. Second, models involving reversible state transitions, such as two-way migration or recombination between loci, include a potentially infinite number of events. Solving the GF for such cases involves matrix inversions (Hobolth *et al.,* 2011; Lohse *et al.,* 2011). Third, while assuming infinite sites mutations may be convenient mathematically and realistic for closely related sequences, this assumption becomes problematic for more distantly related outgroups that are used to polarise mutations in practice. Finally, being able to uniquely map mutations onto genealogical branches assumes phased data that are rarely available for diploid organisms, given the limitations of current sequencing technologies.

In the first part of this paper, we discuss each of these problems in turn and introduce several strategies to remedy the explosion of terms and computation time. These arguments apply generally, irrespective of the peculiarities of particular demographic models and sampling schemes, and suggest a computational “pipeline” for likelihood calculations for non-trivial samples of individuals (up to *n* = 6). The accompanying *Mathematica* notebook implements this scheme for a wide range of demographic histories that may involve arbitrary divergence, admixture and migration between multiple populations, as well as population size changes. As a concrete example, we describe maximum likelihood calculations for a model of isolation with continuous migration (IM) between two populations for unphased and unpolarized data from two diploid individuals. We compare the power of this scheme to that of minimal samples of a single haploid sequence per population. Finally, to illustrate the new method, we estimate divergence and migration between the butterfly species *Heliconius melpomene* and *H. cyndo* (Martin *et al.,* 2013).

## Models and Methods

### Partitioning the GF into equivalence classes

Because the GF is defined in terms of genealogical branches and each topology is specified by a unique set of branches, an intuitive strategy for computing likelihoods is to partition the GF into the contributions from different topologies. To condition on a certain topology, we simply set GF terms that are incompatible with it to 0 (Lohse *et al.,* 2011). Importantly however, such incompatible events still contribute to the total rate Σ*_i_* λ*_i_* of events in the denominator of equation 2. Then, setting all *ω* in the topology-conditioned GF to zero gives the probability of that particular topology. Although conditioning on a particular topology gives a GF with a manageable number of terms, it is clearly not practical to do this for all possible topologies, given their sheer number even for moderate *n* (Table 1).

**Table 1:**
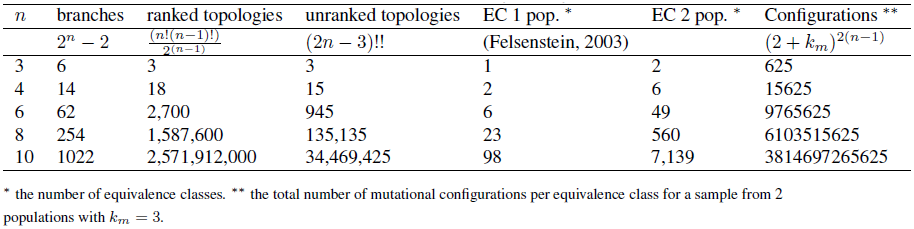
Fundamental quantities of genealogies.

In the following, we will distinguish between ranked and unranked topologies. The GF is a sum over all possible sequences of events in the history of a sample; Edwards (1970) called them “labelled histories”. Considering only coalescence events, each labelled history corresponds to a ranked topology, i.e. a genealogy with unique leaf labels and a known order of nodes. A fundamental property of the standard coalescent, which follows directly from the exchangeability of genes sampled from the same population, is that all ranked topologies are equally likely (Hudson, 1983; Kingman, 1982). In other words, if we could somehow assign each mutation to a particular coalescence (i.e. internode) interval, we could use a much simpler GF, defined in terms of the (n − 1) coalescence intervals rather than the 2(n − 1) branches for inference. This logic underlies demographic methods that use the branch length information contained in well-resolved genealogies (e.g. Nee *et al.,* 1995; Pybus *et al.,* 2002) and coalescent-based derivations of the site frequency spectrum (Griffiths & Tavaré, 1998; Chen, 2012).

Unfortunately however, when analysing sequence data from sexual organisms, we are generally limited by the number of mutations on any one genealogical branch and so often cannot resolve all nodes or their order. Although unranked topologies are not equiprobable, even under the standard coalescent, their leaf labels are still exchangeable. Therefore, each unranked, unlabelled topology, or “tree shape” *sensu* Felsenstein (1978, 2003), is an equivalence class that defines a set of identically distributed genealogies (Fig. 1). This means we only need to work out the GF for one representative (random labelling) per equivalence class. The full GF can then be written as a weighted sum of the GFs for such class representatives:

**Figure 1:**
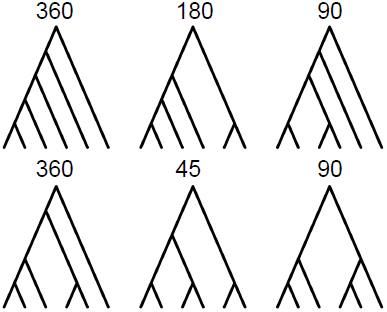
Unranked, unlabelled topologies define equivalence classes of genealogies. For a sample of *n* = 6 from a single population there are six equivalence classes. Their size, i.e. the number of labelled genealogies in each class (*n_h_*) is shown above.

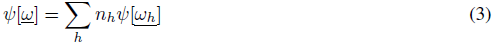
where, *n_h_* denotes the size of equivalence class *h* and *<u>ω_h_</u>* ⊂*<u>ω</u>* is the set of dummy variables that corresponds to the branches of a single class representative in *h*. There are necessarily many fewer equivalence classes than labelled topologies (Table 1). For example, given a sample of size *n* = 6 from a single population, there are 945 unranked topologies, but only six equivalence classes (Fig. 1).

Crucially, the idea of tree shapes as equivalence classes extends to any demographic model and sampling scheme. For samples from multiple populations, the equivalence classes are just the permutations of population labels on (unlabelled) tree shapes. It is straightforward to generate and enumerate the equivalence classes (Felsenstein, 2003) for any sample. For example, for a sample of *n* = 6 from each of two populations (three per population), there are 49 equivalence classes (partially labelled shapes), which can be found by permuting the two population labels on the unlabelled tree shapes in figure 1.

In general, the size of each equivalence class *n_h_* is a function of the number of permutations of individuals on population labels. For *n_i_* individuals from population *i*, there are *n_i_*! permutations. Since the orientation of nodes is irrelevant, each symmetric node (i.e. connected to identical subclades) in the equivalence class halves the number of unique permutations:

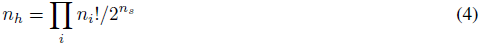

where *n_s_* is the number of symmetric nodes.

Any tree shape contains at least one further symmetry: there is at least one node which connects to two leaves. Because the branches descending from that node have the same length by definition, we can combine mutations (and hence *ω* terms) falling on them: E.g. for a triplet genealogy with topology (*a*, (*b,c*)), we can combine mutations on branch *b* and *c* without loss of information. The joint probability of seeing a configuration with *k_b_* and *k_c_* mutations can be retrieved from *P*[*k_b_* + *k_c_*]:

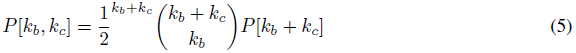

We have previously made use of this in implementing likelihood calculations for triplet samples (Lohse et *al.,* 2011). Although in principle, this combinatorial argument extends to arbitrary genealogies, one can show that, for larger samples, computing *P*[*<u>k</u>*] from mutational configurations defined in terms of internode intervals is computationally wasteful compared to the direct calculation (see File S1).

### Approximating models with reversible events

Migration and recombination events are fundamentally different from coalescence and population divergence. Going backwards in time, they do not lead to simpler sample configurations. Thus, the GF for models involving migration and/or recombination is a system of coupled equations the solution of which involves matrix inversion and higher order polynomials and quickly becomes infeasible for large *n* (Hobolth *et al.,* 2011). As an example, we consider two populations connected by symmetric migration at rate *M = 4Nm.* Given that in practice we are often interested in histories with low or moderate migration, it seems reasonable to consider an approximate model in which the number of migration events is limited. Using a Taylor series expansion, the full GF can be decomposed into histories with 1, 2,… *n* migration events (Lohse et *al.,* 2011). Note that the same argument applies to to recombination between discrete loci and can be used to derive the GF for the sequential Markov coalescent (McVean & Cardin, 2005). It is crucial to distinguish between *M* terms in the numerator and denominator. In other words, even if we stop including sampling configurations involving multiple migration events, *M* still contributes to the total rate Σ*_i_* λ*_i_* in the denominator. We can modify the GF for a pair of genes *a* and *b* sampled from two populations connected by symmetric migration (Lohse et *al.,* 2011, eq. 9) to include an indicator variable γ that counts the number of migration events:

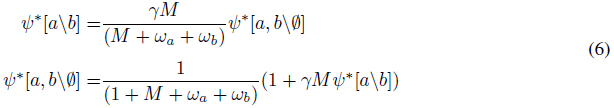

Expanding *ψ\** in γ, the coefficients of γ, γ^2^… γ*^n^* correspond to histories with 1, 2,… *n* migration events. This is analogous to conditioning on a particular topology: the truncated GF does not sum to one (if we set the *ω* to zero), but rather gives the total probability of seeing no more than *n_max_* events. This is convenient in practice because it immediately gives an estimate of the accuracy of the approximation. Expanding the solution of equation 6 around γ = 0 gives:

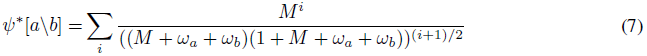

The GF conditional on there being at most one migration event is

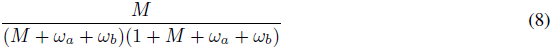

The error of this approximation is:

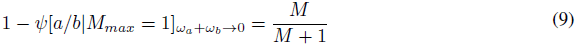
which is just the chance that a migration event occurs before coalescence (see Fig. 2). An analogous expansion for the pairwise GF for the IM model (Lohse *et al.,* 2011, eq. 13) gives:

**Figure 2:**
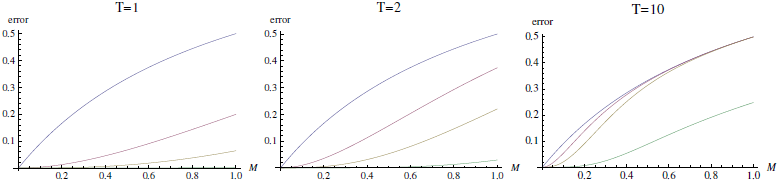
The error in limiting the number of migration events to *n_max_* = 1 (eqs. 1 & 2) (red), 2 (yellow) and 4 (green) for a pairwise sample in the IM model plotted against *M* for different divergence times *T*. The results for a model of equilibrium migration without divergence is shown for comparison (blue).

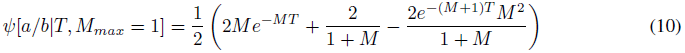

Expressions for the GF conditional on a maximum of 2,3,… *n* migration events and for larger samples can be found by automating the GF recursion. While these do not appear to have a simple form, plotting the error against *M* and *T* (Fig. 2), shows that for recent divergence (*T < 1*) and moderate gene flow (*M < 0.5*), histories involving more than two migration events are extremely unlikely (*p* < 0.01) and can be ignored to a good approximation. Considering that for large n, coalescence, which occurs at rate *n*(*n* − 1)/2, becomes much more likely than migration (at rate *Mn*), this approximation should be relatively robust to sample size.

### Unknown phase and root

There are at least two further complications for block-wise likelihood computations in practice: First, the direct correspondence between mutation types and genealogical branches we have assumed so far, assumes that the infinite sites mutation model holds between in and outgroup, which is often unrealistic in practice. Second, given the current limitations of short read sequencing technology, genomic data are often unphased and one would ideally incorporate phase ambiguity explicitly rather than ignore it (e.g. Lohse & Frantz, 2014) or rely on computational phasing.

Both unknown phase and root can be incorporated via a simple relabeling of branches. In generating the GF, we have labelled branches and corresponding *ω* variables by the tips (leaf-nodes) they are connected to. Crucially, the full GF expressed as a sum over equivalence class representatives has unique labels for all individuals, i.e. we distinguish genes from the same population. To incorporate unknown phase, we simply combine branches with the same set of descendants in each population. Each branch combination correspond to an entry in the (joint) site frequency spectrum (SFS). Consider for example two genes from each of two populations. There are six equivalence classes of rooted genealogies (Fig. 3). Combining all branches with the same population labels gives seven *ω* variables that correspond to unphased site types: *ω_a_*, *ω_b_ ω_ab_ ω_aa_*, *ω_bb_*, *ω_aab_*, *ω_abb_*. In the absence of root information, we further combine the two branches on either side of the root. Denoting *ω* variables for unrooted branches by * and the two sets of individuals they are connected to we have: 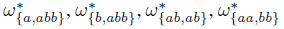. The rooted branches contributing to each unrooted branch are indicated in colour in figure 3. The *ω\** terms correspond to the four types of variable sites defined by the (folded) SFS for two populations: 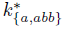 (heterozygous sites unique to *a*), 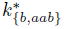 (heterozygous sites unique to *b*), 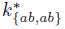 (heterozygous sites shared by both) and 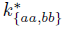 (fixed differences between *a* and *b*). Note also that without the root, the six equivalence classes collapse to two unrooted equivalence classes (defined by branches 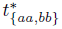 and 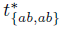) (Fig. 3).

**Figure 3:**
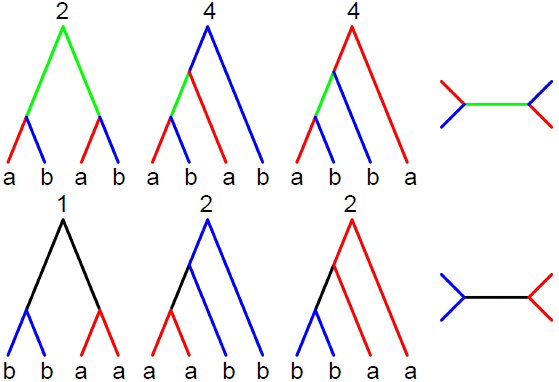
For a sample of two sequences from each of two populations (*a* and *b*), there are six classes of equivalent, rooted genealogies (left); their sizes *n_h_* are shown above. Without root information, these collapse to two unrooted genealogies (right). Without phase information, there are four mutation types that map to specific branches in the rooted genealogy: heterozygous sites unique to one sample 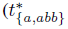 and 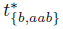, red and blue respectively), shared heterozygous sites (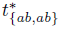, green) and fixed, homozygous differences (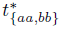, black).

**Figure 4:**
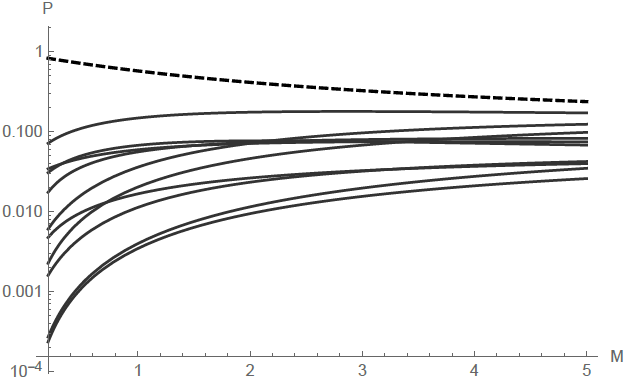
The topology spectrum for a sample of *n* = 6 from a two population IM model with asymmetric migration and *T* = 1.5. The probabilities of all 11 unrooted topolgies are plotted against *M*. The probability of the most likely topology of reciprocal monophyly (((*a,* (*a, a*)), (*b,* (*b, b*))) is shown as a dashed line.

The combinatorial arguments outlined above extend to arbitrary sample sizes and numbers of populations. We modify eq. 9 to write the GF of an unrooted genealogy *ψ*[*<u>ω*</u>*] as a sum over unrooted equivalence classes (denoted *h\**), each of which is in turn a sum over rooted equivalence classes:

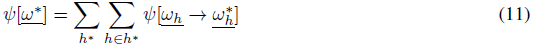

Similarly, the GF for unphased data is given by combining *ω* variables with the same number of descendants in each population. From this simplified GF, we can compute the probability of blockwise counts of mutation types defined by the SFS. Following Bunnefeld *et al.* (2015), who have used this extension of the SFS to block-wise data to fit bottleneck histories in a single population, we will refer to it as the blockwise site frequency spectrum (bSFS).

### Limiting the total number of mutational configurations

In principle, we can compute the probability of seeing arbitrarily many mutations on a particular branch from equation 1. In practice however, the extra information gained by explicitly including configurations with large numbers of mutations (which are very unlikely for short blocks) is limited, while the computational cost increases. An obvious strategy is to tabulate exact probabilities only up to a certain maximum number of mutations *k_m_* per branch and combine residual probabilities for configurations involving more than *k_m_* mutations on one or multiple branches. As described by Lohse *et al.* (2011) and Lohse *et al.* (2012), the residual probability of seeing more than *k_m_* mutations on a particular branch s is given by

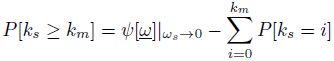

i.e. we subtract the sum of exact probabilties for configurations involving up to *k_m_* mutations from the marginal probability of seeing branch *s*.

Assuming that we want to distinguish between all 2 (*n* − 1) branches in a given equivalence class and use a global *k_m_* for all branches, there are (*k_m_* + 2) possible mutation counts per branch (including those with no mutations or more than *k_m_* mutations on a branch) which gives (*k_m_* + 2)^2(n−1)^ mutational configurations in total. For example, for *n* = 6 and *k_m_* = 3 there are 9,765,625 mutational configurations per equivalence class (Table 1). Although this may seem daunting, most of these configurations are extremely unlikely, so a substantial computational saving can be made by choosing branch-specific *k_m_*. We have implemented functions in *Mathematica* to tabulate *P*[*<u>k</u>*] for an arbitrary vector of *k_m_* (File S1).

The bSFS with *k_m_* = 0 defines mutational configurations by the joint presence and/or absence of mutation types defined by the SFS in a block, irrespective of the number of mutations of each type. This constitutes an interesting special case. In the limit of very large blocks, i.e. if we assume an unlimited supply of mutations, this converges to the topological probabilities of equivalence classes which can be obtained directly from the partitioned GF by setting all *ω* → 0. We can think of this set of probabilities as the “topology spectrum”. For a sample of 3 genes from each of 2 populations this consists of 49 equivalence classes which reduce to 11 unrooted topologies (Fig. 6). Under the IM model with unidirectional migration, the GF of each class is solvable using *Mathematica* (see Supplementary.nb). The most likely topology is reciprocal monophyly, i.e. (((*a, a*)*, a*)), ((*b, b*)*, b*)))). As expected, its probability decreases with *M* and increases with *T*.

## Results

The various strategies for simplifying likelihood calculations based on the GF outlined above suggest a general “pipeline”, each component of which can be automated:

1. Generate all equivalence classes *h* and enumerate their sizes *n_h_* for a given sampling scheme.
2. Generate and solve the GF conditional on one representative within each *h*.
3. Take the Inverse Laplace Transform with respect to the parameters that correspond to discrete events (e.g. divergence, admixture, bottlenecks). These processes are initially modelled as occurring with a continuous rate.
4. Re-label *ω* variables to combine branches and equivalence classes that are indistinguishable in the absence of root and/or phase information.
5. Find sensible *k_m_* cut-offs for each mutation type from the data.
6. Tabulate probabilities for all mutational configurations in each equivalence class.

In the accompanying *Mathematica* notebook we have implemented this pipeline as a set of general functions. These can be used to automatically generate, solve and simplify the GF (step 1–3 above), and – from this – tabulate *P*[*<u>k</u>*], the likelihood of a large range of demographic models (involving population divergence, admixture and bottlenecks) (6 above). In principle, this automation works for arbitrary sample sizes. In practice however, the inversion step (3 above) and the tabulation of probabilities (6 above) become prohibitively slow for *n* > 6.

To give a concrete example, we derive the GF for a model of isolation (at time *T* × 2*N_e_* generations) with migration (at rate *M* = 4*N_e_m* migrants per generation) (IM) between two populations (labelled *a* and *b*). We further assume that migration is unidirectional, i.e. from *a* to *b* forwards in time and that both populations and their common ancestral population are of the same effective size (we later relax this assumption when analysing data). As above, we consider the special case of a single diploid sample per population without root and phase information. We first derive some basic properties of unrooted genealogies under this model. We then investigate the power of likelihood calculations based on the bSFS. Finally, we apply this likelihood calculation to an example dataset from two species of *Heliconius* butterflies.

### The distribution of unrooted branches under the IM model

We can find the expected length of any branch (or combination of branches) *s* from the GF as: 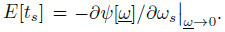 The expressions for the expected lengths of rooted branches are cumbersome (File S2). Surprisingly however, the expected lengths of the four unrooted branches 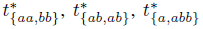 and 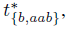 each of which is a sum over the underlying rooted branches (Fig. 3), have a relatively simple form (Fig. 5):

**Figure 5:**
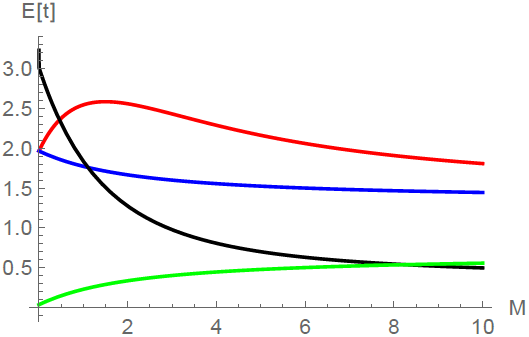
The expected length of unrooted genealogical branches (eq. 12) for a sample of *n* = 4 under the IM model of two populations (*a* and *b*) with asymmetric migration and population divergence time T = 1.5 (×2*N_e_* generations). Colours correspond to those in figure 3.

**Figure 6:**
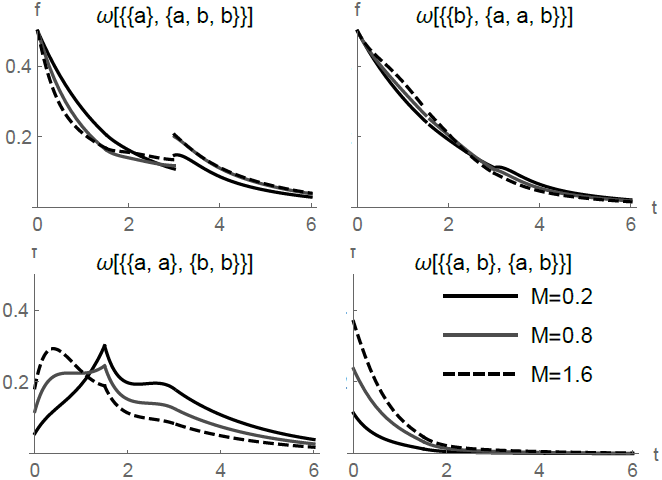
The length distribution of unrooted genealogical branches for a sample of *n* = 4 under the IM model of two populations (*a* and *b*) with asymmetric migration and population divergence at *T* = 1.5 (in 2*N_e_* generations).

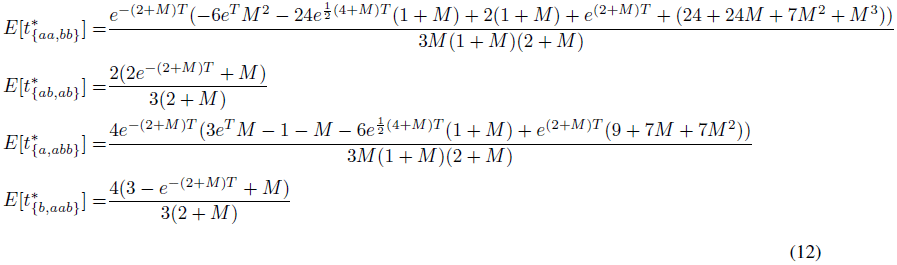

Similarly, the probability of the two unrooted topologies reduces to:

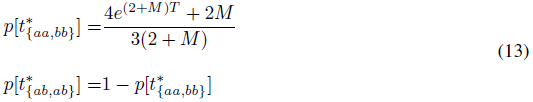

We can recover the full distribution of rooted branches from the GF by taking the Inverse Laplace Transform (using *Mathematica*) with respect to the corresponding *ω*\*. While this does not yield simple expressions (File S2), examining figure 6 illustrates that much of the information about population history is contained in the shape of the branch length distribution rather than its expectation (Fig. 5). For example, branches carrying fixed differences 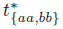 have a multi-modal distribution with discontinuities at *T* and the relative size of the first mode depends strongly on *M*.

### Power analysis

We compared the power to detect post-divergence gene flow between two different blockwise likelihood calculations: the bSFS for a diploid genome per population (*n* = 4) and a minimal sample of a single haploid sequence (*n* = 2) per population. We measured power as the expected difference in support (*E*[Δ*lnL*]) between the IM model and a null model of strict divergence without gene flow and arbitrarily assumed datasets of 100 blocks. However, since we are assuming that blocks are unlinked, i.e. statistically independent, (*E*[Δ*lnL*]) scales linearly with the number of blocks.

Figure 7 shows the power to detect gene flow for a relatively old split (T = 1.5) and sampling blocks with an average of 1.5 heterozygous sites within each species (i.e. *θ = 4N_e_μ* = 1.5). Without gene flow, this corresponds to a total number of 5.2 mutations per block on average. Unsurprisingly, sampling a diploid sequence from each population gives greater power to detect gene flow than pairwise samples (compare black and blue lines in figure 7). However, contrasting this with the power of a simpler likelihood calculation for *n* = 4 which is based only on the total number of mutations *S_T_* in each block (grey line in figure 7), illustrates that the additional information does not stem from the increase in sample size *per se,* but rather the addition of topology information. In fact, there is less information in a larger sample without topology information than in pairwise samples. Similarly, adding root information almost doubles power (green lines in Fig. 7).

**Figure 7:**
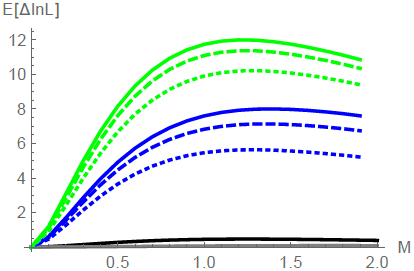
The power (*E*[Δ*lnL*]) to distinguish between an IM model and a null model of strict divergence (*T* = 1.5) from 100 unlinked blocks of length *θ* = 1.5 for different sample sizes and data summaries: the total number of mutations in a sample of *n* = 2 (black) and *n* = 4 (grey), the bSFS for unphased data for two diploids (*n* = 4) with root (green) and without root (blue). Dotted, dashed and solid lines correspond to different maximum numbers of mutations per branch type, *k_m_* = 0, 1 and 3 respectively.

In comparison and perhaps surprisingly, the threshold *k_m_* has relatively little effect on power. In other words, for realistically short blocks, most of the information is contained in the joint presence and absence of mutation types (regardless of their number), i.e. *k_m_* = 0.

### *Heliconius* analysis

To illustrate likelihood calculation based on the bSFS, we estimated divergence and gene flow between two species of *Heliconius* butterflies. The sister species *H. cydno* and *H. melpomene rosina* occur in sympatry in parts of Central and South America, are known to hybridise in the wild at a low rate (Mallet *et al.,* 2007), and have previously been shown to have experienced post-divergence gene flow (Martin *et al.,* 2013). We sampled 150 bp blocks of intergenic, autosomal sequence for one individual genome of each species from the area of sympatry in Panama (chi565 and ro2071). These data are part of a larger resequencing study involving high coverage genomes for four individuals of each *H. cydno* and *H. m. rosina* as well as an allopatric population of *H. melpomene* from French Guiana (Martin *et al.,* 2013). We excluded CpG islands and sites with low quality (GQ <30 and MQ<30), excessively low (<10) or high (>200) coverage and only considered sites that passed these filtering criteria in all individuals.

We partitioned the intergenic sequence into blocks of 225bp length and sampled the first 150 bases passing filtering in each block. 6.3% of blocks violated the 4-gamete criterion (i.e. contained both fixed differences and shared heterozygous sites) and were removed. This sampling strategy yielded 161,726 blocks with an average per site heterozygosity of 0.017 and 0.015 in *H. m. rosina* and *H. cydno* respectively (Fig. 8). Summarizing the data by counting the four mutation types in each block gave a total of 2,337 unique mutational configurations, 1,743 of which occured more than once.

**Figure 8:**
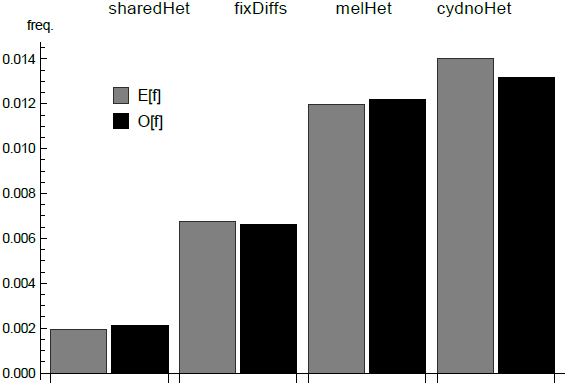
The folded SFS has four site types: i) heterozygous sites unique to either *H. m. melpomene* or ii) *H. cydno* iii) shared in both species and iv) fixed differences. The observed genome wide SFS is shown in black. The expectation under the IM history estimated from the bSFS (Table 2) was computed using eq. 12 and is shown in grey.

**Table 2:** Top: Support (Δ*lnL* relative to the best model) for isolation with migration and strict divergence (Div) between *H. m. rosina* and *H. cydno.* Migration from *H. cydno* into *H. m. rosina* (*IM_c→m_*) fits better than migration in the opposite direction (*IM_m→c_*). Bottom: Maximum likelihood estimates of parameters under the *IM_c→m_* model (scaled estimates in brackets).

| *Div* | $IM_{m \rightarrow c}$ | $IM_{c \rightarrow m}$ | | |
| --- | --- | --- | --- | --- |
| -49.1 | -26.2 | 0 |  |  |
| $\theta (N_e)$ | $\theta_C (N_e)$ | $T$ | $M$ | |
| 1.25 ($1.10 \times 10^6$) | 3.24 ($2.85 \times 10^6$) | 1.90 (1.04 MY) | 1.50 | |

We initially used all blocks (regardless of linkage) to obtain point estimates of parameters under three models: i) strict isolation without migration (*Div*) ii) isolation with migration from *H. cydno* into *H. m. rosina* (*IM_c→m_*) and iii) isolation with migration from *H. m. rosina* into *H. cydno* (*IM_c→m_*). In all cases, we assumed that the common ancestral population shared its *N_e_* only with one descendant species while the other species has a different *N_e_*. We maximise *lnL* under each model using Nelder-Mead simplex optimisation implemented in the *Mathematica* function *NMaximise.* To compare models, we corrected for LD by rescaling Δ*lnL* with a factor of 1/121. This admittedly *ad hoc* correction was obtained after examining the decay of LD between pairs of blocks with distance (for scaffolds >200kb) (File S2). At a distance of 121 blocks (which corresponds to an average physical distance of >27kb) the correlation drops below 0.025 and LD approaches background levels (see Discussion).

We find strong support for a model of isolation with migration from *H. cydno* into *H. m. rosina* (*IM_c→m_*) (Table 2). This model fits significantly better than both a history of strict divergence or divergence followed by migration in the opposite direction (*IM_m→c_*). Our results agree with earlier genomic analyses of these species that showed support for post-divergence gene flow based on D-statistics (Martin *et al.,* 2013), IMa analyses based on smaller numbers of loci (Kronforst *et al.,* 2013) and genome wide SNP frequencies analysed using approximate Bayesian computation. Asymmetrical migration from *H. cydno* into *H. m. rosina* has also been reported previously, and could be explained by the fact that F1 hybrids resemble *H. m. rosina* more closely due to dominance relationships among wing patterning alleles, possibly making F1s more attractive to *H. melpomene* (Kronforst *et al.,* 2006; Martin *et al.,* 2015).

A recent direct, genome-wide estimate of the mutation rate for *H. melpomene* (Keightley *et al.,* 2015) allows us to convert parameter estimates into absolute values. Assuming a spontaneous mutation rate of 29 × 10^−9^ per site and generation and using the ratio of divergence between *H. m. rosina* and the more distantly-related ‘silvaniform’ clade of *Heliconius* at synonymous and intergenic sites to estimate selective constraint on intergenic sites, gives an effective mutation rate of *μ* = 1.9 × 10^−9^ (Martin *et al.,* 2015). Applying this rate to our estimate of *θ* and assuming four generations per year, we obtain an *N_e_* estimate of 1.10 × 10^6^ for *H. m. rosina* and the common ancestral population and 2.85 × 10^6^ for *H. cydno.* We estimate species divergence to have occurred roughly 1 million years ago. Note that this is more recent than previous estimates of 1.5 million years which was obtained using approximate Bayesian computation and a different calibration based on mitochondrial genealogies (Kronforst *et al.,* 2013; Martin *et al.,* 2015).

## Discussion

We have shown how the probabilities of genealogies, and hence of mutational configurations, can be calculated for a wide variety of demographic models. This gives an efficient way to infer demography from whole genome data. Irrespective of any particular demographic history, the possible genealogies of a sample can be partitioned into a set of equivalence classes, which are given by permuting population labels on tree shapes. We show how this fundamental symmetry of the coalescent can be exploited when computing likelihoods from blockwise mutational configurations. We have implemented this combinatorial partitioning in *Mathematica* to automatically generate and solve the generating function (GF) of the genealogy and, from this, compute likelihoods for a wide range range of demographic models. Given a particular sample of genomes, we first generate a set of equivalence classes of genealogies and condition the recursion for the GF (Lohse *et al.,* 2011) on a single representative from each class. This combinatorial strategy brings a huge computational saving. Importantly, it does not sacrifice any information. This is in contrast to a similar partitioning of the GF, which as we show, can be used to find approximations for models that include reversible events, in particular migration between populations and recombination between discrete loci and involves a trade-off between computational efficiency and loss of information.

Although these approaches make it possible to solve the GF for surprisingly large samples and biologically interesting models, the number of mutational configurations (which explodes with the number of sampled genomes) remains a fundamental limitation of likelihood calculations in practice. Given outgroup and phase, the full information is contained in a vast table of mutational configurations which are defined in terms of the 2(*n* − 1) branches of each equivalence class. For samples from two populations, the number of mutational configurations we need to calculate is the product of the last two columns of Table 1. For example, considering a sample of 3 haploid genomes per populations and allowing for up to *k_m_* = 3 mutations per branch, there are 49 × 9,765,625 = 478, 515,625 possible mutational configurations.

### The bockwise site frequency spectrum

Our initial motivation for studying the bSFS was to deal with unphased data in practice. The GF of the bSFS can be obtained from the full GF simply by combining branches with equivalent leaf labels. As well as being a lossless summary of blockwise data in the absence of phase information, the bSFS is a promising summary in general for several reasons. First, it is extremely compact compared to the full set of (phased) mutational configurations. Unlike the latter, the size of the bSFS does not depend on the number of equivalence classes (which explodes with *n*, Table 1), but only on *n*. Given a sample of *n_i_* individuals from population *i* and assuming a global maximum number of mutations *k_m_* for all mutation types, the (unfolded) bSFS comprises of a maximum of ((Π*_i_*(*n_i_* + 1)) − 2)^(^*^km^*^+2)^) mutational configurations. For a sample of 3 haploid genomes from each of two populations and *k_m_* = 3, the bSFS has 7^5^ = 16,807 entries. Second, because equivalence classes of genealogies are defined by the presence and absence of SFS types, much of the topology information contained in the full data will still be captured in the bSFS. Finally, and perhaps surprisingly, at least for the IM model the expressions for the total length of branches contributing to unphased and unpolarized mutation types (eq. 12 & 13) are much simpler than those of the underlying rooted branches, which suggests that it may be possible to find general results.

Despite the strategies developed here, it is clear that full likelihood calculations will rarely be feasible for samples > 6 given the rapid increase in the number of equivalence classes. However, a separation of timescales exists for many models of geographic and genetic structure (Wakeley, 1998, 2009), and so full likelihood solutions for moderate (*n* < 6) samples may be sufficient for computing likelihoods for much larger samples if these contribute mainly very short branches with no mutations in the initial scattering phase during which lineages from the same population either coalesce or trace back to unsampled demes.

### Dealing with linkage

A key assumption of our blockwise likelihood calculations is that there is no recombination within sequence blocks, and that different blocks are independent of each other. This latter assumption is especially problematic when we analyse whole-genome data. If we divide the genome into blocks that are small enough for recombination within them to be negligible, our method correctly gives the probabilities of possible mutational configurations, and this can be used to fit a demographic model. However, the accuracy of this fit will be grossly overestimated if we simply multiply likelihoods across blocks, because adjacent blocks are strongly correlated. Ignoring this correlation is essentially a composite likelihood calculation. Suppose, that we multiply likelihoods across every *k* th block, *k* being chosen large enough that blocks are uncorrelated. This procedure is valid starting at any block, and so can be repeated *k* times, such that the whole genome is included in the analysis, and taking the average *lnL* across all *k* analyses. This is equivalent to simply multiplying the likelihoods across all blocks, and then dividing the total *lnL* by *k*. In the *Heliconius* example above, we found that there is little correlation between blocks 121 blocks apart, and so assessed significance simply by dividing the *lnL* by 121. We note that analysing well-separated blocks or SNPs is very common practice (e.g. Wang & Hey, 2010; Excoffier *et al.,* 2013), and is essentially equivalent to our simple procedure. However, this procedure is quite arbitrary, and clearly needs improvement. On the one hand, successive blocks or SNPs are not completely correlated, suggesting that this procedure considerably underestimates the accuracy of estimates. On the other hand, however, there may be weak, long-range correlations, due to a small fraction of long regions that coalesced recently, and these may increase the variance of parameter estimates. The safest course is to check the accuracy of estimates by simulation under the inferred demographic model and a realistic model of recombination via a full parametric bootstrap.

An advantage of direct likelihood calculations is that one can easily check the absolute fit of the data to a model by asking how well the observed frequency of mutational configurations or some summary such as the SFS is predicted by the model. For example, the IM history we estimated for the two *Heliconius* species fits the observed genome-wide SFS reasonably well (Fig. 8). The fact that we slightly underestimate the heterozygosity in *H. cydno* may suggests that some process (e.g. demographic change after divergence or admixture from an unsampled ghost population/species) is not captured by our model.

In general, the GF framework makes it possible to derive the distribution of any summary statistic that can be defined as a combination of genealogical branches and understand its properties under simple demographic models and small *n*. Although explicit calculations based on such summaries are not feasible for large *n*, summary statistics such as the bSFS may still have wide applicability for fitting complex models and larger samples of individuals, for example using approximate likelihood methods, or simply as a way to visualize how genealogies vary along the genome.

## Acknowledgements

This work was supported by funding from the UK Natural Environment Research Council to KL (NE/I020288/1) and a grant from the European Research Council (250152) to NB. We thank Lynsey Bunnefeld for helpful discussion and comments.

## Data Availability

File S1 is a *Mathematica* notebook that contains the code to generate the GF and tabulate likelihoods under arbitrary demographic models. File S2 contains the code used for the analyses of the IM model, including the analyses of the *Heliconius* data and the power test. The processed input data for *Heliconius* and python scripts used are available from www.datadryad.com doi:XXX; raw sequence data are published by Martin *et al.* (2013) and available from www.datadryad.com doi:10.5061/dryad.dk712.

